# Comammox *Nitrospira* bacteria outnumber canonical nitrifiers irrespective of nitrogen source and availability

**DOI:** 10.1101/2021.09.27.461883

**Authors:** Katherine J. Vilardi, Irmarie Cotto, Maria Sevillano Rivera, Zihan Dai, Christopher L. Anderson, Ameet Pinto

## Abstract

Complete ammonia oxidizing bacteria coexist with canonical ammonia and nitrite oxidizing bacteria in a wide range of environments. Whether this is due to competitive or cooperative interactions, or a result of niche separation is not yet clear. Understanding the factors driving coexistence of nitrifiers is critical to manage nitrification processes occurring in engineered and natural ecosystems. In this study, microcosm-based experiments were used to investigate the impact of nitrogen source and loading on the population dynamics of nitrifiers in drinking water biofilter media. Shotgun sequencing of DNA followed by co-assembly and reconstruction of metagenome assembled genomes revealed clade A2 comammox bacteria were likely the primary nitrifiers within microcosms and increased in abundance over *Nitrsomonas-like* ammonia and *Nitrospira-like* nitrite oxidizing bacteria irrespective of nitrogen source type or loading. Changes in comammox bacterial abundance did not correlate with either ammonia or nitrite oxidizing bacterial abundance in urea amended systems where metabolic reconstruction indicated potential for cross feeding between ammonia and nitrite oxidizing bacteria. In contrast, comammox bacterial abundance demonstrated a negative correlation with nitrite oxidizers in ammonia amended systems. This suggests potentially weaker synergistic relationships between ammonia and nitrite oxidizers might enable comammox bacteria to displace nitrite oxidizers from complex nitrifying communities.

## Introduction

Nitrification, the biological transformation of ammonia to nitrate via nitrite, is an important process in engineered and natural ecosystems. While nitrification mediated by ammonia oxidizing microorganisms (AOM) (Kowalchuk & Stephen, 2001, Stahl & de la Torre, 2012), including ammonia oxidizing bacteria (AOB) and archaea (AOA), and nitrite oxidizing bacteria (NOB) (Daims *et al*., 2016) has been extensively investigated, complete ammonia oxidation (comammox) performed by comammox bacteria is understudied in large part due to its recent discovery. All known comammox bacteria belong to *Nitrospira* sub-lineage II (Daims *et al*., 2015, van Kessel *et al*., 2015, Pinto *et al*., 2016) and are currently divided into two clades, A and B, with clade A further separated into sub-clades A1 and A2 (Palomo *et al*., 2019). Due to close phylogenetic relatedness, comammox-*Nitrospira* cannot be distinguished from *Nitrospira*-NOB based on the 16S rRNA gene sequence or the marker genes for nitrite oxidation (*nxrAB*) (Daims *et al*., 2015). Thus, characterization of comammox bacteria has been largely enabled by shotgun DNA sequencing followed by reconstruction of assembled genomes (Palomo *et al*., 2016, Pinto *et al*., 2016, Camejo *et al*., 2017, Wang *et al*., 2017, Annavajhala *et al*., 2018, Poghosyan *et al*., 2019) and the development of primers targeting subunits of comammox bacteria ammonia monooxygenase (*amo*) gene (Bartelme *et al*., 2017, Pjevac *et al*., 2017, Fowler *et al*., 2018, Wang *et al*., 2018, Beach & Noguera, 2019, Cotto *et al*., 2020).

Within the engineered water cycle, clade A1 comammox bacteria have been primarily detected in wastewater treatment plants while clade A2 and B have been associated with drinking water treatment and distribution systems (Palomo *et al*., 2019). It is unclear if this translates into physiological differences between the clades/sub-clades since there is only one comammox isolate and an enrichment whose kinetic parametes have been reported. To date, kinetic parameters of comammox bacteria are confined to two clade A representatives, cultured *Candidatus* N. inopinata and an enrichment of *Candidatus* N. kreftii (Kits *et al*., 2017, Sakoula *et al*., 2020). Both demonstrate a high affinity for ammonia, with half-saturation constants orders of magnitude lower than strict AOB. Comparatively, the *Candidatus* N. kreffti enrichment exhibited a higher affinity for nitrite compared to *Candidatus* N. inopinata and demonstrated partial inhibition of ammonia oxidation even at low ammonia concentrations (Sakoula *et al*., 2020). This suggests that clade-specific comammox bacterial niche, if applicable, may be arise from a combination of factors ranging from affinity to inhibition. Beyond clade specific traits, identifying the potential environmental and physiological factors driving the coexistence of comammox bacteria with canonical nitrifiers is also important to better understand comammox bacteria role in complex nitrifying communities (Gulay *et al*., 2019, Liu *et al*., 2019, Wang *et al*., 2019, Wang *et al*., 2019, Zheng *et al*., 2019, Gottshall *et al*., 2020, Wang *et al*., 2020, He *et al*., 2021, Shao & Wu, 2021). Comammox bacteria have been detected along with their canonical nitrifying counterparts in wastewater treatment plants (Gonzalez-Martinez *et al*., 2016, Roots *et al*., 2019, Zheng *et al*., 2019, Cotto *et al*., 2020, Yang *et al*., 2020), drinking water systems (Pinto *et al*., 2016, Tatari *et al*., 2017, Wang *et al*., 2017, Fowler *et al*., 2018, Poghosyan *et al*., 2020) and soils (Prosser & Nicol, 2012, Shi *et al*., 2018, Liu *et al*., 2019, He *et al*., 2021) at varying abundances over a wide range of ammonium concentrations. While there is currently no quantitative estimate of the contribution of comammox bacteria to nitrification compared to AOB and NOB, several studies have investigated comammox bacterial dynamics in the context of mixed nitrifying communities. For instance, DNA/RNA stable isotope probing provided support for comammox *Nitrospira* contributing to ammonia oxidation in lab-scale biofilters exposed to very low ammonium concentrations (Gulay *et al*., 2019). Soil microcosms amended with high ammonia concentrations were enriched in AOB compared to those with lower ammonia concentrations where clade B comammox bacteria proliferated (Wang *et al*., 2019, He *et al*., 2021). Interestingly, in a lab-scale partial nitrification-anammox reactor operating with incrementally increased ammonia loadings, comammox bacteria initially dominated over strict AOB but its abundance significantly declined as loadings were further increased (Shao & Wu, 2021).

Comammox bacteria may also acquire ammonia via urea degradation. Specifically, genes encoding for urea transport and the urease enzyme are distributed among many *Nitrospira* populations (Koch *et al*., 2015), including most comammox populations (Palomo *et al*., 2018). While this may diversify potential nitrogen sources for comammox bacteria (Daims *et al*., 2016), this could be a potential advantage for canonical nitrifiers involved in a reciprocal feeding strategy as observed with co-cultured *Nitrospira moscoviensis* converting urea to ammonia for *Nitrosomonas europaea* (Koch *et al*., 2015). The tight interplay between canonical nitrifiers is well established; however, our understanding of comammox competition (or lack thereof) with AOM and its impact on strict NOB in mixed communities is limited.

To better understand the comammox bacterial role within these complex nitrifying communities, we investigated their population dynamics across two nitrogen sources (ammonia or urea) at three total nitrogen dosing strategies. Thus, the objectives of this study were (1) to determine if comammox bacteria and canonical nitrifiers exhibit concentration and nitrogen source dependent dynamics when subject to repeat nitrogen amendments and (2) to determine if these dynamics are consistent or variable at the clade or population within each functional guild. Characterization of microbial communities in biofilters at drinking water treatment plants has revealed rich nitrifier diversity (Fowler *et al*., 2018, Gulay *et al*., 2019), making it an ideal sample source for this study. Collectively, our microcosm-based study offers novel insights regarding the ecophysiology of clade 2 associated comammox bacteria; information on this clade are very limited. Further, while other microcosm studies have focused on competitive interactions between comammox bacteria and strict AOB under controlled conditions (Wang *et al*., 2019, He *et al*., 2021), there is only limited assessment of NOB response to experimental treatment. This study explicitly assesses the NOB dynamics in response to nitrogen source and loading rates in the context of the broader nitrifying community.

## Materials and Methods

### Experimental design and execution

Granular activated carbon (GAC) with coexisting AOB, NOB, and comammox bacterial populations from biofilters at the drinking water treatment plant (DWTP) in Ann Arbor, (AA) Michigan was used as the inoculum for this experimental work (Pinto *et al*., 2016). Microcosms consisted of 3 grams of GAC supplemented with 10 mL of filter influent from AA DWTP in 40 mL pre-sterilized glass vials (DWK Life Sciences – Fisher 033395C). A total of 96 glass microcosms were prepared such that two biological replicates for each of the three nitrogen concentrations (1.5, 3.5 and 14 mg-N/L) for the two nitrogen sources (i.e., ammonium (direct) and urea (indirect)) were harvested weekly for analyses over the period of the 8-week experiment. Ammonium was spiked in at 0.1, 0.25 and 1 mM (in the form of ammonium chloride solution), corresponding to final concentrations of 1.5, 3.5 and 14 mg-N/L. For urea, 0.05, 0.125 and 0.5 mM (in the form of urea solution) were used to ensure similar concentrations of total nitrogen as the ammonium microcosms. Microcosms were maintained by removing approximately 10 mL of spent filter influent and subsequently replenishing them with 10 mL of fresh influent and the respective nitrogen source spike every two days. Once a week, two microcosms per condition (i.e., nitrogen concentration and nitrogen source) were sacrificed and two 0.5 g GAC samples from each microcosm were transferred to Lysing Matrix E tubes (MP Biomedical Lysing Matrix E – Fisher MP116914100) and stored at −80°C until further processing. Additionally, aqueous samples were collected and filtered through 0.2 μM filters (Sartorius Minisart NML Syringe Filter – Fisher Scientific 14555269) for chemical analyses. The sampled aqueous volume was replaced with fresh substrate to increase the ammonia and urea concentrations to microcosm specific concentrations. Hach Company Test n’ Tube Vials were used to determine concentrations of ammonia-N (Hach, Cat No. 2606945), nitrite-N (Hach, Cat No. 2608345) and nitrate-N (Hach, Cat No. 2605345) in microcosms. All samples were analyzed on a Hach DR1900 photospectrometer (Hach – DR1900-01H). Alkalinity of filtered liquid samples were measured using Hach Alkalinity Total TNTplus Vials (Hach – TNT870).

### DNA extraction and qPCR

GAC samples were subjected to DNA extraction using the DNAeasy PowerSoil kit (Qiagen, Inc – Cat No.12888) on the QIAcube (Qiagen, Inc – Cat No. 9002160) following manufacturer’s instructions with a few modifications. Specifically, the lysing buffer from the PowerBead tubes were transferred to the Lysing Matrix E tubes and C1 buffer was added. Prior to bead beating, an equal volume of chloroform was added (610 μL). Bead beating consisted of four rounds of 40 seconds on a FastPrep-24 instrument (MP Bio – 116005500) with bead beading tubes placed on ice for two minutes between each bead beating. Samples were then centrifuged at 10,000 g for 1 minute and 750 μL of aqueous phase used to purify DNA using the QIAcube Protocol for the DNeasy PowerSoil Kit. Each round of extractions included a reagent blank as a negative control. After extraction, DNA concentration was determined using a Qubit instrument with the dsDNA Broad Range Assay (ThermoFisher Scientific – Cat No. Q32850) (Table S1). DNA was stored in a −80<C freezer until future use.

qPCR assays were conducted using QuantStudio 3 Real-Time PCR System (ThermoFisher Scientific – Cat. No. A28567). Primer sets targeting the 16S rRNA gene of AOB (Hermansson & Lindgren, 2001), 16S rRNA gene of *Nitrospira* (Graham *et al*., 2007), *amoB* gene of clade A comammox bacteria (Cotto *et al*., 2020) and 16S rRNA gene for total bacteria (Caporaso *et al*., 2011) were used (Table S2). Previously published primer set for the *amoB* gene of clade A comammox bacteria was updated based on metagenomic data generated as part of this study (Cotto *et al*., 2020). Based on alignments of *amoB* gene sequences from the comammox MAGs assembled in this study, the previously published forward primer for comammox clade A *amoB* from Cotto et. al 2020 had one mismatch with one of our bins. Thus, this forward primer was further modified by changing the 13^th^ position from G to a degenerate base S (seven base pairs from 3’-end). The use of the modified primers resulted in increased abundance of comammox bacteria in this study as shown in supplementary Figure S1, indicating the ability to capture comammox *amoB* gene sequences not amplified by previous primer set.

The qPCR reactions were carried out in 20 μL volumes, which included 10 μL Luna Universal qPCR Master Mix (New England Biolabs, Inc., Cat. No. NC1276266), 5 μL of 10-fold diluted template DNA, primer concentrations are outlined in Table S4 and DNAse/RNAse free water (Fisher Scientific, Cat. No. 10977015) to make up the remaining volume to 20 μL. Each sample per assay was subject to qPCR in triplicate and qPCR plates were prepared using the epMotion M5073 liquid handling system (Eppendorf, Cat. No. 5073000205D). The cycling conditions used in this study were as follows: initial denaturing at 95°C for 1 minute, 40 cycles of denaturing at 95°C for 15 seconds, annealing temperatures and time used are listed in Table S2 and extension at 72°C for 1 minute. qPCR analysis proceeded with a negative control and 7-point standard curve ranging from 10^3^-10^9^ copies of 16S rRNA gene of *Nitrosomonas europaea* for total bacteria quantification, 10^2^-10^8^ copies of 16S rRNA genes of *Nitrosomonas europaea* and *Ca* Nitrospira inopinata for AOB and *Nitrospira* quantification, respectively, and 10^2^ – 10^8^ copies of *amoB* gene of *Ca* Nitrospira inopinata for the quantification of comammox bacteria. The primer used to detect the 16S rRNA gene of *Nitrospira* would inclusively track both comammox-*Nitrospira* and *Nitrospira*-NOB. Thus, *Nitrospira*-NOB abundance was estimated by subtracting the copy number of comammox bacteria *amoB* from the copy number of 16S rRNA gene of *Nitrospira*.

### Metagenomic analyses

A subset of samples were selected for metagenomic analysis including DNA extracted from the initial GAC inoculum and samples from weeks four and eight (n=13) for all nitrogen sources and dosing strategies. DNA extracts from duplicate microcosms for each time point were pooled in equal mass proportion before sending DNA templates for sequencing at the Roy J. Carver Biotechnology Center at University of Illinois Urbana-Champaign Sequencing Core. Two lanes of Illumina NovaSeq were used to generate paired-end reads ranging from 29 to 68 million per sample (2×150-bp read length) (Table S3). Raw paired-end reads were trimmed and quality filtered with fastp (Chen *et al*., 2018) (Table S5). Filtered reads were mapped to the UniVec Database (National Center for Biotechnology Information) using BWA (Li & Durbin, 2009) to remove potential vector contamination. Subsequent unmapped reads were extracted, sorted and indexed using SAMtools v1.3.1 (Li *et al*., 2009) then converted back to FASTQ using bedtools v2.19.1 (Quinlan & Hall, 2010).

Small subunit rRNA sequence reconstruction from quality filtered short reads was carried out using the Phyloflash v3.4 (Gruber-Vodicka *et al*., 2020). Briefly, bbmap was used to map short reads against the SILVA 138.1 NR99 database with the default minimum identity of 70% followed by assembly of full-length sequences with Spades (kmers = 99,111,127) and detection of closest-matching database sequences using usearch global within VSEARCH at a minimum identity of 70%. For read pairs, taxonomic classification was performed by taking the lowest common ancestor using SILVA taxonomy (Pruesse *et al*., 2007). Assembled sequences from all samples belonging to nitrifying bacteria were clustered at 99% identity using vsearch v2.15.2 (Rognes *et al*., 2016). Reference *Nitrospira* and *Nitrosomonadaceae* 16S rRNA reference sequences were obtained from ARB-SILVA and aligned with assembled sequences using muscle v3.8.1551 (Edgar, 2004). Construction of 16S rRNA phylogenetic trees for *Nitrospira* and *Nitrosomonadaceae* was performed using IQ-TREE v1.6.12 (Nguyen *et al*., 2015) with model finder option (Kalyaanamoorthy *et al*., 2017) selecting TIM3+F+I+G4 and TPM2u+F+I+G4 as models for respective trees.

Quality filtered paired-end reads from all samples were co-assembled with metaSPAdes v3.11.1 (Nurk *et al*., 2017) with k-mers lengths 21, 33, 55, 77, 99, and 119, and phred off-set of 33. Quality evaluation of the assembled scaffolds was performed using Quast v5.0.2 (Gurevich *et al*., 2013) (Table S4). Open reading frames (ORF) on scaffolds were predicted using Prodigal v2.6.2 (Hyatt *et al*., 2010) with the “meta” flag and functional prediction of resulting protein sequences were determined by similarity searches of the KEGG database (Hiroyuki Ogata, 1999) using kofamscan (Aramaki *et al*., 2020). Taxonomic classification of scaffolds harboring nitrogen cycling genes was performed using kaiju v1.7.4 (Menzel *et al*., 2016) against the NCBI nr database with default parameters. CoverM v0.5.0 (www.github.com/wwood/CoverM) was used to calculate reads per kilobase million (RPKM) of these scaffolds as a metric for estimating relative abundance in each sample.

Scaffolds were binned into clusters and manually refined using Anvi’o (v5.1 and 5.5) (Eren *et al*., 2015) with three binning algorithms including CONCOCT (Alneberg *et al*., 2014), Metabat2 v2.5 (Kang *et al*., 2019) and Maxbin2 v2.2.7 (Wu *et al*., 2016). DAS_tool v1.1.2 (Sieber *et al*., 2018) was used to merge bins from the three approaches to generate final metagenome assembled genomes (MAGs). Completeness and contamination of the final set was determined using CheckM v1.0.7 (Parks *et al*., 2015) followed by taxonomic classification using the Genome Taxonomy Database Toolkit v1.2.0 with release 89 v04-RS89 (Chaumeil *et al*., 2019). CoverM was used to calculate RPKM for each bin. Similar to the annotation of the metagenome, functional prediction of bin ORFs were determined by similarity searches against the KEGG database using kofamscan. The annotation of genes of interest were further confirmed by querying protein sequences against the NCBI-nr database using BLASTP. MAGs were also annotated using Prokka as a secondary annotation method (Seemann, 2014). The Up-to-date Bacterial Core Gene pipeline (UBCG) (Na *et al*., 2018) with default parameters was used to extract and align a set of 92 singe copy core genes from *Nitrospira* and *Nitrosomonas* references genomes (Table S5) and nitrifier MAGs for phylogenomic tree reconstruction. Maximum likelihood trees were generated based on the nucleotide alignment using IQ-TREE with model finder selecting the GTR+F+R10 and GTR+F+R4 models for *Nitrospira* and *Nitrosomonas* trees, respectively, with 1000 bootstrap iterations. For outgroups, two *Leptospirillum* and three *Nitrosospira* genomes were used for *Nitrospira* and *Nitrosomonas* trees, respectively. Pairwise alignments of comammox *amoA* and *hao* and Nitrospira *nxrA* protein sequences were created using muscle. Maximum likelihood trees were inferred by IQ-TREE with model finder selecting LG+G4 for the *amoA* tree and LG+I+G4 for *hao* and *nxrA* trees with 1000 bootstrap iterations for each tree. The *amoA* and *hao* protein sequences from *Nitrosomonas europaea* and *Nitrosomonas oligotropha* were used as the outgroup for comammox trees. All trees were visualized using the Interactive Tree of Life (itol) (Letunic & Bork, 2019). Pairwise comparisons of average nucleotide identity of 38 *Nitrospira* and 15 *Nitrosomonadaceae* genomes (Table S5) with nitrifier MAGs obtained in this study was determined using FastANI v1.31 (Jain *et al*., 2018).

### Statistical analysis

The relative abundance of each nitrifier population was tested to determine if significant differences existed between concentration or source of electron donor types using ANOVA and Welch t-tests, respectively, with R version 4.0.4. Shapiro Wilks tests were used to confirm normality prior to these statistical tests. Linear regression and correlation analysis were used to examine the relationship between the abundance of nitrifying guilds in each of the nitrogen amendments over time.

## Results

### Microbial community composition in microcosms and nitrogen biotransformation potential

Microcosms consisting of granular activated carbon (GAC) from drinking water biofilters were subject to intermittent amendments of nitrogen using two nitrogen sources (ammonia or Urea) across three nitrogen concentrations (14, 3.5, and 1.5 mg-N/L). The conditions used in these experiments are denoted as 14A, 3.5A, 1.5A, 14U, 3.5U and 1.5U where A or U represents ammonia or urea amendments, respectively, and the number represents the concentration of nitrogen source spike in mg/L as nitrogen. Two microcosms were sacrificed on a weekly basis over the duration of a eight week experiment (n=96 total microcosms). Extracted DNA from the inocula and weeks four and eight were subject to shotgun DNA sequencing (n=13).

Initial assessment of taxonomic diversity in the samples based on analyses of metagenomic reads mapping to the small subunit rRNA database (SILVA SSU NR99 version 138.1) indicated that the GAC inocula largely consisted of bacteria with archaea and eukaryota constituting a small proportion of the overall metagenomic reads (~0.002%). The bacterial community was primarily composed of Gammaproteobacteria (20-30%), Alphaproteobacteria (25-31%) and Nitrospirota (8-15%) (Figure 1A). *Nitrospira* and *Nitrosomonadaceae* were the only nitrifiers identified and constituted 9-15% of the overall microbial community in samples. Full length 16S rRNA gene sequences were assembled from each sample (n=13) resulting in a total of eight sequences with closest matching SILVA database hits to uncultured *Nitrospira* bacteria (Accession numbers: MF040566, AY328760, JN868922). Clustering of all eight *Nitrospira* 16S rRNA gene sequences at 99% identity resulted in two *Nitrospira* operational taxonomic units (OTUs) with one cluster composed of six sequences (Nitrospira OTU 1) and the other cluster with two sequences (Nitrospira OTU 2). Phylogenetic placements of these OTUs revealed both clustered within *Nitrospira* sublineage II (supplementary figure S2A). Diversity of *Nitrospira* was likely underrepresented as full length *Nitrospira* 16S rRNA gene sequences could not be assembled from some samples despite a large portion of extracted 16S rRNA gene reads mapping to *Nitrospira* references in the SILVA database. Limited assembly of these reads could be due to several closely related *Nitrospira* species/strains coexisting in the samples making reconstruction of full length sequences difficult. For canonical AOB, Nitrosomonas sp. AL212 (CP002552) was the closest matching database hit to one assembled sequence while another six had hits closet to *Nitrosomonadaceae* (Accession numbers: FPLP01009519, KJ807851, FPLK01002446) but could not be further classified at the genus or species level. Phylogenetic placement of the single Nitrosomonas OTU affiliated it with *Nitrosomonas* sp. AL212 and *Nitrosomonas ureae* (Figure S2B).

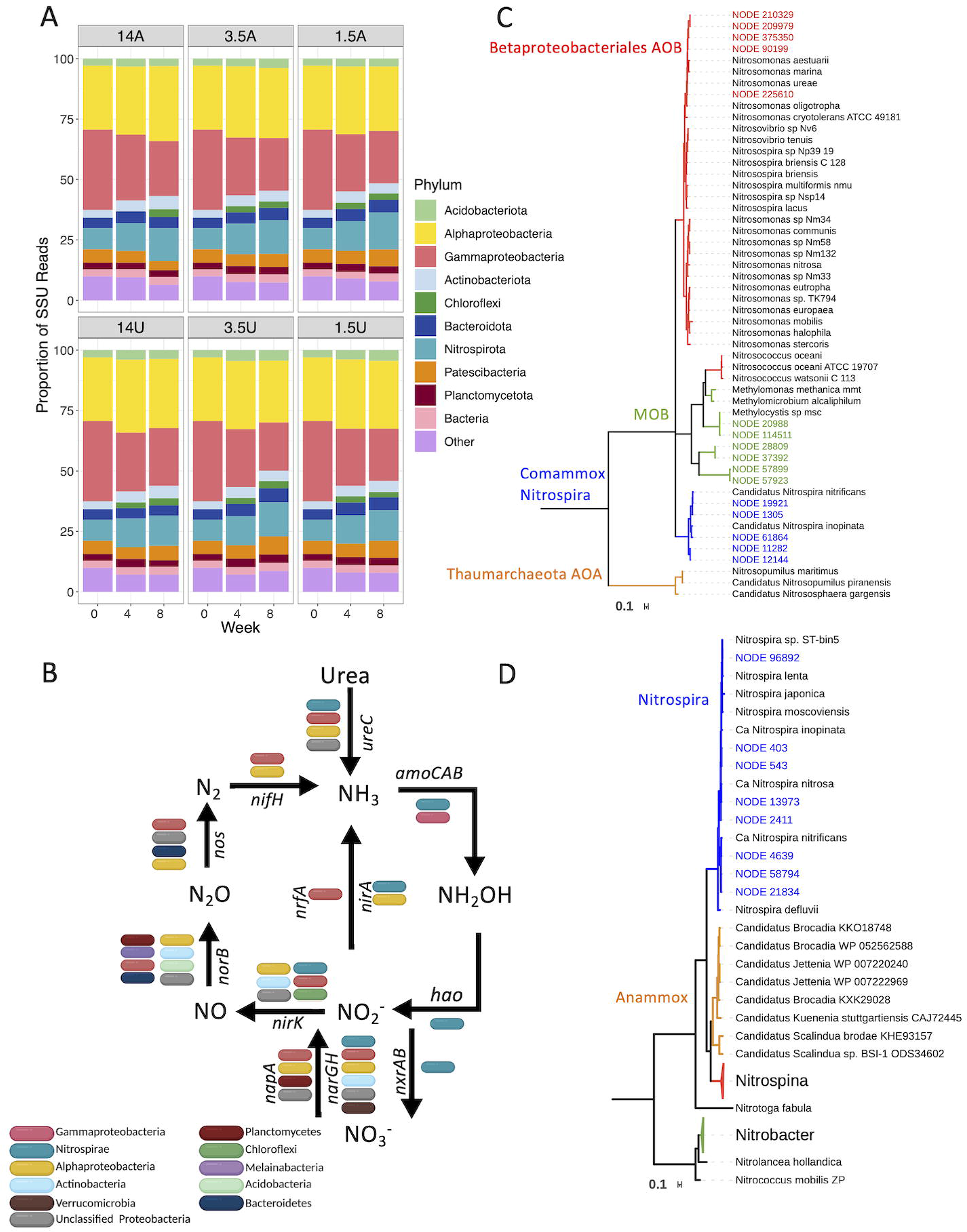

Following co-assembly of metagenomic reads, predicted protein coding genes from scaffolds associated with the nitrogen metabolism were taxonomically classified (Figure 1B). The majority of methane/ammonia monooxygenase (*pmo-amo*) like genes (KEGG orthology: K10944, K10945, K10946) were associated with either nitrifiers (i.e., *Nitrospira* or *Nitrosomonas*) or methanotrophs (i.e., *Methylocystis*) (Figure 1C). While some *amoCAB* genes could not be classified to the genus level using kaiju software, blastp searches against the NCBI non-redundant protein database indicated these were closely related to *Nitrosomonas*. All retrieved *hao* sequences (KEGG orthology: K10535) were associated with Nitrospira which is likely due to the low relative abundance of *Nitrosomonas*-like populations and the resulting inability to assemble their *hao* genes. Potential for ureolytic activity was detected across four phyla based on the urease alpha subunit (*ureC). ureC* sequences associated with Nitrospirota and Gammaproteobacteria could be classified at the genus level as *Nitrospira* and *Nitrosomonas*. Sequences identified as nitrate reductase/nitrite oxidoreductase alpha and beta subunits (K00370, K00371) were subject to further classification to differentiate between nitrite oxidoreductase genes belonging to NOB from nitrate reductases belonging to other community members. Phylogenetic placement of most *Nitrospira nxrA* sequences found in this study cluster within a branch containing both comammox and *Nitrospira*-NOB species (Candidatus *N. inopinata*, Candidatus *N. nitrosa* and *N. defluvii*) (Figure 1D). While other sequences clustered on a separate branch with Candidatus *N. nitrificans*, a single *Nitrospira nxrA* sequence clustered closely within a branch containing only *Nitrospira*-NOB belonging to sublineage II.

### Phylogenomic placement of nitrifying populations and their metabolism

Metagenome assembled genomes (MAGs) were obtained from GAC microcosms after dereplication of MAGs from three binning approaches. All 204 MAGs were classified as bacteria, with 145 MAGs exhibiting completeness greater than 70% and contamination less than 10% (Table S6). Approximately 62% of the metagenomic reads mapped to these MAGs. Nine MAGs in total were classified as nitrifying bacteria belonging to *Nitrosomonas* and *Nitrospira* (Table S7). Genome annotation confirmed that four *Nitrospira* MAGs had key ammonia (ammonia monooxygenase and hydroxylamine oxidoreductase) and nitrite (nitrite oxidoreductase) oxidation genes (Figure S3). Quality assessment for these comammox MAGs indicated two high (Bin_49_2_2 and Bin_49_4) and one medium quality (Bin_260) (Table S1) according to (Bowers *et al*., 2017). A fourth comammox MAG (Bin_13) was assembled with high completeness (89%) but also possessed high redundancy (18%) that could not be improved with further manual refinement. The remaining two *Nitrospira* MAGs (Bin_7_1 and Bin_188), which were likely strict NOB due to lack of ammonia oxidation genes, were less complete (38.04% and 48.25%) with low redundancy (8.76% and 8.46%). The low completeness was likely not due to their lower abundance, but potentially high level of strain heterogeneity which may have affected the assembly of reads associated with *Nitrospira*-NOB. For example, RPKM-based relative abundance estimated using all reads (total RPKM) showed the two *Nitrospira*-NOB MAGs exhibited similar relative abundance to comammox bacteria MAGs Bin_49_2_2 and Bin_49_4 (~7-10 total RPKM), but the CheckM estimated strain heterogeneities for Bin_7_1 and Bin_188 were 40 and 75, respectively, compared to 0 for both Bin_49_2_2 and Bin_49_4. Two MAGs classifying as *Nitrosomonas* were deemed high (Bin_83) and medium quality (Bin_168); however, a third *Nitrosomonas* MAG was considered low quality.

A maximum likelihood tree based on 91 single copy core genes confirmed all *Nitrospira* MAGs affiliated with sublineage II (Figure 2A). Four of the *Nitrospira* MAGs from this study clustered within clade A comammox *Nitrospira* (Bin_49_2_2, Bin_49_4, Bin_260 and Bin_13) but were separated into distinct groups on the phylogenomic tree; namely, forming three clusters with MAGs obtained from tap water, drinking water filters, and freshwater. *amoA*-based phylogenetic analysis corroborated their placement into clade A (Figure 2B); however, *hao-based* phylogeny distinguished three of comammox MAGs (Bin_49_2_2, Bin_49_4, Bin_260) as clade A2 (Palomo et al. 2019) while one clustered within clade A1 (Bin_13) (Figure 2C). Consistent across all trees, Bin_49_2_2 and Bin_260 cluster closely with comammox MAGs *Nitrospira* sp. SG-bin2 and ST-bin4 (ANI ~ 92%) derived from tap water metagenomes (Wang *et al*., 2017). Bin_49_4 clustered closely with Nitrospirae bacterium Ga0074138 (ANI ~ 99%), which was previously detected in GAC from the same drinking water treatment plant (Pinto *et al*., 2016), along with other tap water and groundwater-fed rapid sand filter MAGs (Palomo *et al*., 2016, Wang *et al*., 2017). Bin_13 associated with comammox MAGs obtained from freshwater, UBA5698 and UBA5702 (Parks *et al*., 2017) (ANI ~ 90%); however, its high contamination (18%) likely renders ANI comparison less accurate. Overall, the MAGs demonstrated less then 95% ANI to other reference comammox bacterial MAGs (Figure S4) suggesting comammox bacteria detected in GAC microcosms are distinct from one another and previously published comammox MAGs; as a result, they are likely novel *Nitrospira* species. The two remaining *Nitrospira* MAGs, Bin_7_1 and Bin_188, clustered with strict *Nitrospira*-NOB MAGs recovered from tap water, Nitrospira_sp_ST-bin5 (Wang *et al*., 2017) (ANI ~ 94%), and a rapid sand filter, Nitrospira CG24D (ANI ~ 87%) (Palomo *et al*., 2016) (Figure 2A and S3). However, since Bin_7_1 and Bin_188 were highly incomplete, the possibly they are novel comammox bacteria cannot be excluded. Only two strict AOB MAGs (Bin_83 and Bin_168) from this study were used for phylogenomic analysis due high redundancy and low completeness of the third (Bin_195). Both Bin_83 and Bin_168 originate from *Nitrosomonas* cluster 6a and clustered closely with *Nitrosomonas ureae* and *Nitrosomonas* sp. AL212 (Figure 2D). Bin_168 shares a high sequence similarity to *N. ureae* (ANI ~ 98%) while Bin_83 shares less than 83% ANI to any of the references on the tree including Bin_168.

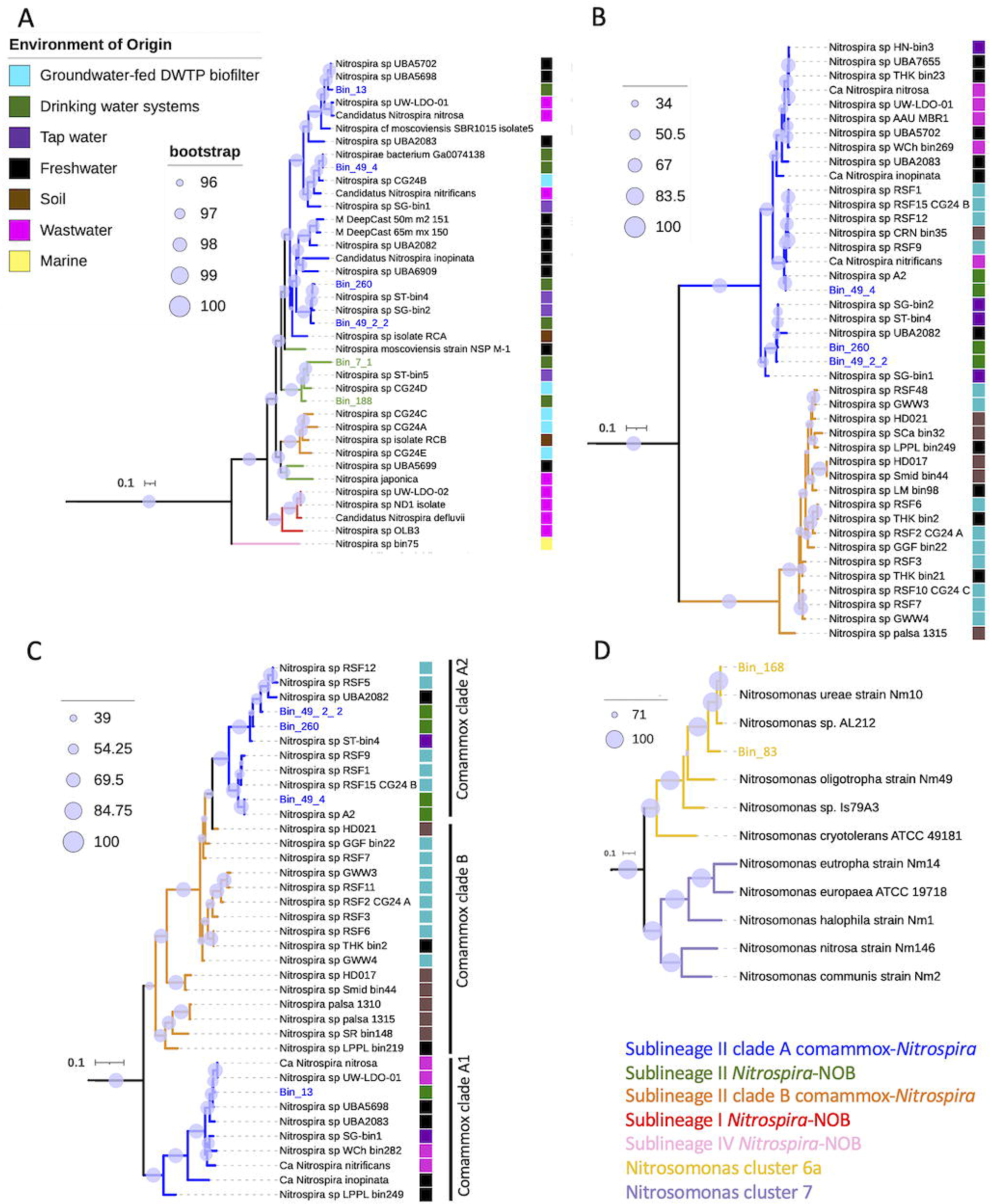

All comammox MAGs demonstrated the potential for ureolytic activity with the presence of the *ureABC* operon in addition to most genes for urease accessory proteins (Figure S2). *Nitrospira*-NOB MAGs did not contain genes encoding for urease; however, two *ureC* sequences found on assembled scaffolds that were classified as *Nitrospira* but were not binned into any of the Nitrospira MAGs. Queries of these *ureC* genes against the NCBI non-redundant database revealed one sequence shared the highest percent identity to *Nitrospira lenta* and *Nitrospira moscoviensis* while top hits for the second sequence belonged to an unclassified *Nitrospira*. One *Nitrospira*-NOB MAG (Bin_7_1) did harbor genes for the urea transport system permease proteins (*urtBC*), urea transport system substrate-binding proteins (*urtA*) and urea transport system ATP-binding proteins (*urtDE*). This suggests that the two unbinned *ureC* genes likely belonged to *Nitrospira*-like NOB bacteria. *Nitrosomonas* MAGs Bin_168 and Bin_83 each contained the *ureCAB* operon and some genes for urease accessory proteins and urea transport. A third *ureC* sequence found in the metagenome classified as *Nitrosomonas* but was not binned into any *Nitrosomonas* MAGs.

### The impact of nitrogen amendments on nitrifying populations

To address concentration and nitrogen source-dependent dynamics of the three nitrifier populations detected in our metagenomic analysis, qPCR-assays were used to estimate their abundances over time in the nitrogen amended microcosms. In the high ammonia amendment (14A), strict AOB relative abundance increased 2.4-fold from weeks 1-3 but remained below 2% of total bacteria for the duration of the experiment whereas comammox relative abundance increased markedly over time reaching 2.8% of total bacteria by end of the experiment (Figure 3B). Similar to strict AOB, *Nitrospira*-NOB relative abundance increased early on but thereafter reduced from 4% at its peak in week two to 1.8% by week eight. Weekly measurements for nitrogen concentrations taken alongside biomass samples indicated the presence of residual ammonia and accumulated nitrite concentrations were highest during the first three weeks of the experiment but gradually reduced over time with most inorganic nitrogen present as nitrate (Figure S5). While comammox bacteria were always dominant, qPCR-based abundance of strict AOB as a portion of AOM was significantly higher when ammonia and nitrite accumulated in weeks 1-3 as compared to weeks 5-8 (Welch’s t-test, p-value < 0.05) (Figure 3A).

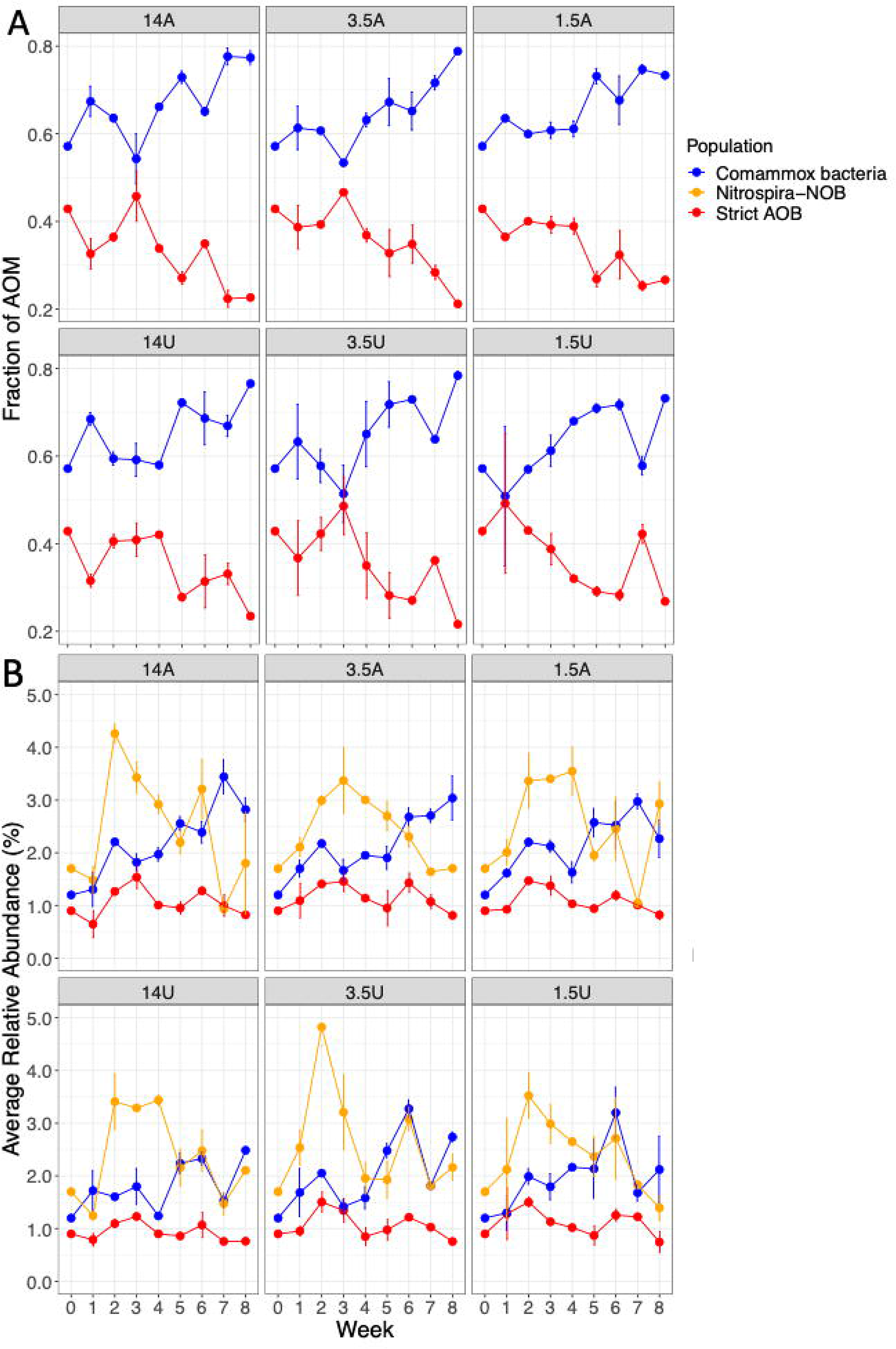

The qPCR data was in concordance with metagenomic estimation of MAG abundance with clade A2 comammox (Bin_49_2_2, Bin_49_2 and Bin_260) highly abundant compared to strict AOB (Bin_83, Bin_168 and Bin_195) and clade A1 comammox (Bin_13) in the inocula and at weeks four and eight in the high ammonia amendment (Figure 4). In particular, clade A2 MAGs Bin_49_2_2 and Bin_49_4 were the most dominant comammox populations while strict AOB was dominated by Bin_83 at each time point. *Nitrospira*-NOB MAGs had comparable abundance to clade A2 comammox MAGs but displayed limited variation in abundance in the high ammonia amendments. This contrasts with the qPCR data, where *Nitrospira*-NOB were significantly more abundant than comammox bacteria at earlier timepoints and then demonstrated a significant decrease in abundance over time. This is likely due to the fact that the two assembled Nitrospira-NOB MAG’s do not represent the entirety of NOB diversity in the microcosms as several *nxr* genes were not binned into MAGs and that metagenomic data is only available for select timepoints as compared to qPCR data.

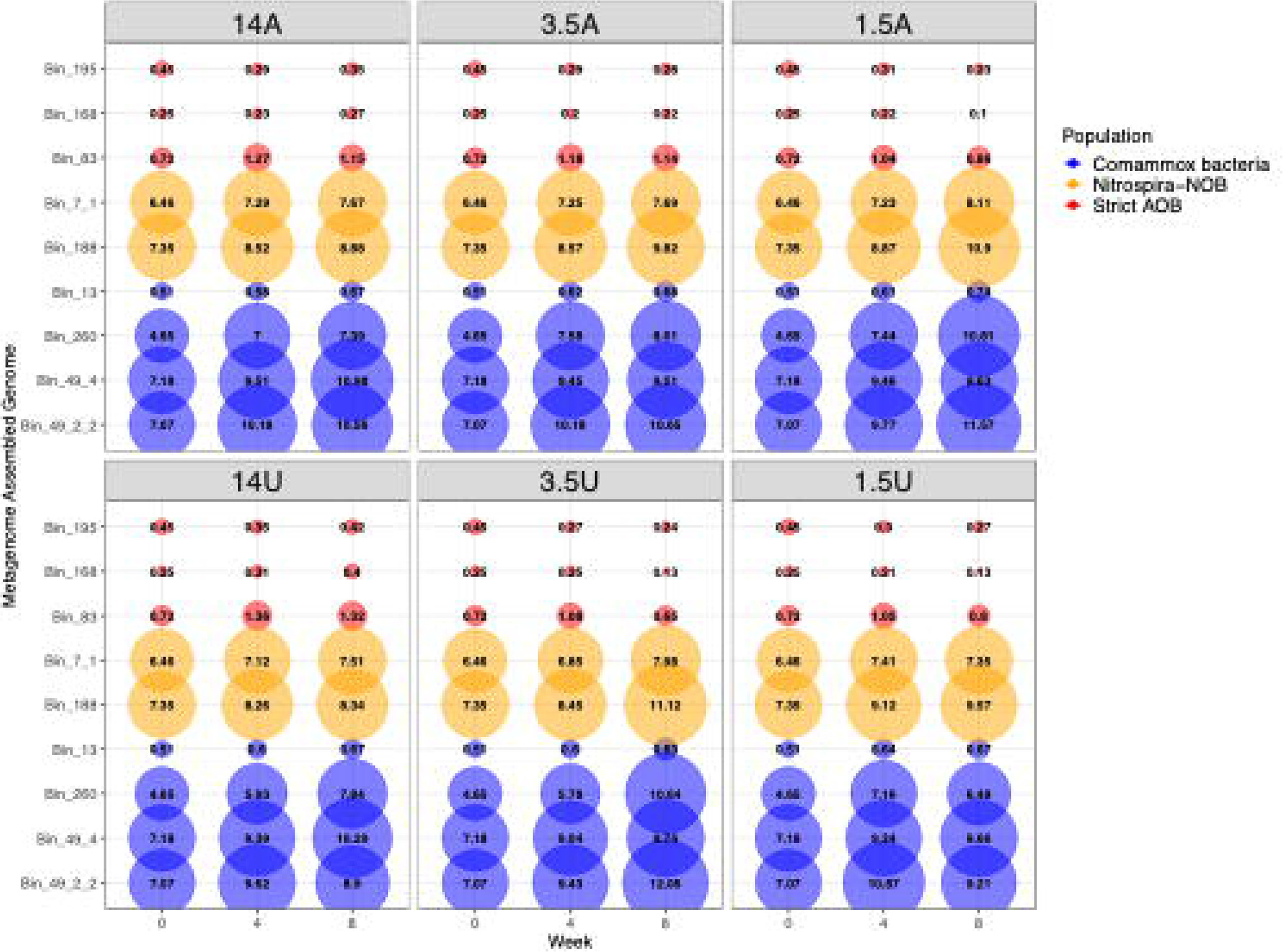

Nitrifier populations in mid and low ammonia amendments displayed similar dynamics to those observed in high ammonia with comammox relative abundance increasing to 3% and 2.2% of total bacteria by week eight, respectively. Interestingly, Bin_260, the least abundant clade A2 comammox MAG in the inocula, demonstrated significant increase in abundance in the low ammonia amendment over the course of the experiment compared to its abundance in the other ammonia amendments. Consistent with the ammonia amended microcosms, strict AOB in urea amended microcosms increased in relative abundance only at earlier time points followed by low but stable relative abundance (~2% of total bacteria). In the high urea amendment, relative abundance of comammox bacteria remained largely unchanged at earlier time points followed by an increase in abundance. Despite this, mean relative abundance of comammox bacteria compared to strict AOB was still approximately 2-fold greater in all urea amendments. Similar to the ammonia amendments, *Nitrospira*-NOB relative abundance did increase initially followed by a decline in all urea amendments. Interestingly though, the relative abundance of comammox bacteria and *Nitrospira*-NOB were similar in the later weeks of the experiment after *Nitrospira*-NOB’s initial rise in urea amendments. Clade A2 comammox MAG Bin_260 was consistently lower in abundance than Bin_49_2_2 and Bin_49_2 in the urea amendments except for mid urea. Abundance of the clade A1 comammox MAG remained lower than all clade A2 MAGs and displayed minimal enrichment in all the urea amendments which was consistent with ammonia amended microcosms. Bin_168, which showed high sequence similarity to *Nitrosomonas ureae*, did not exhibit enrichment in any of the urea amendments and remained low in abundance with all other strict AOB MAGs.

There was no significant difference in the mean qPCR-based relative abundance of strict AOB or *Nitrospira*-NOB between the high ammonia (14A) and urea amendments (14U) (Welch t-test, p > 0.05) but the mean relative abundance of comammox bacteria was significantly greater in high ammonia than in the high urea amendment (Welch t-test, p < 0.05). Comparatively, out of all nitrogen amendments, mean relative abundance of comammox bacteria was the lowest in high urea (1.8% of total bacteria). Comparisons between the mid ammonia (3.5A) and urea amendments (3.5U) as well as the low ammonia (1.5A) and urea (1.5U) amendments revealed no significant difference in mean relative abundance for any of the nitrifier populations (Welch t-test, p > 0.05). Additionally, no significant differences were detected when testing the mean relative abundance of the three nitrifier populations between high, mid, and low concentrations within each amendment type (ANOVA, p > 0.05).

The relative abundance of the nitrifying groups were used to examine potential correlations between the different populations in each of the nitrogen amendments. The ratio of comammox bacteria as portion of AOM to comammox bacteria as a portion of total *Nitrospira* revealed a strong positive relationship in all amendments (Pearson R = 0.75-0.87, p < 0.001) (Figure S6A), however, the change in relative abundance of comammox bacteria was not directly correlated with that of strict AOB in any of the nitrogen amendments (Figure S6B). Strict AOB and *Nitrospira*-NOB abundances were strongly correlated for all urea amendments and high (14A) and mid ammonia (3.5A) (Pearson r = 0.58-0.82, p < 0.05, Figure 5A) but exhibited a weaker relationship in low ammonia (Pearson r = 0.42, p > 0.05). Interestingly, while comammox bacteria abundance was significantly and negatively correlated with that of *Nitrospira*-NOB in ammonia amendments (Pearson r = −0.37 to −0.61) (p < 0.05), there was no significant association between them in the urea amendments (p > 0.05) (Figure 5B).

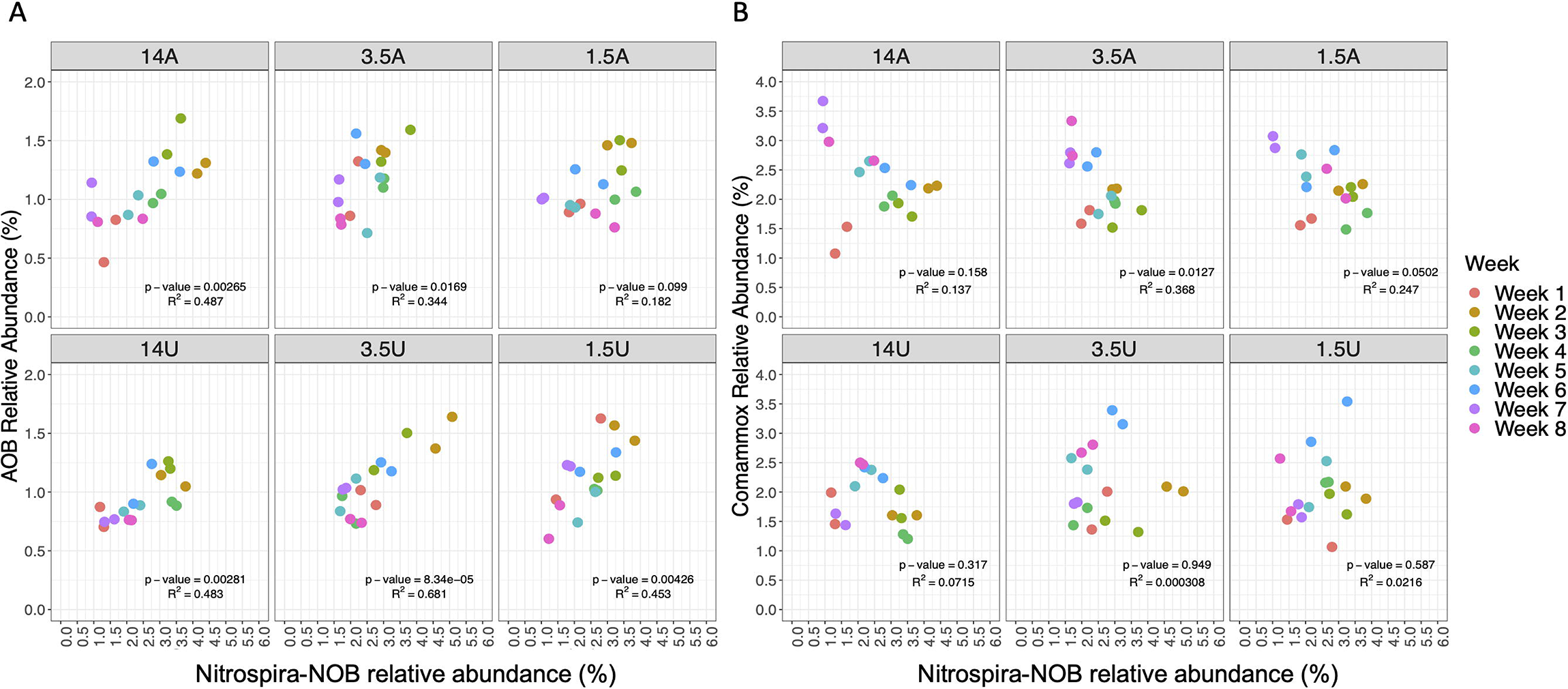

## Discussion

### Key nitrifiers encompassing Nitrospira and Nitrosomonas-like bacteria share ureolytic potential

16S rRNA gene sequences assembled from short reads indicated *Nitrospira*- and *Nitrosomonas*-like populations were the only nitrifiers present in the microcosms. The proportion of 16S rRNA gene reads mapping to *Nitrospira*-like populations in this study suggested that they were highly abundant in the inocula and nitrogen amendments. Surveys of other DWTP biofilters using 16S rRNA gene amplicon sequencing have indicated that sublineage II *Nitrospira* account for a dominant portion of the bacterial community (Gulay *et al*., 2016) with further investigation confirming high contributions to its abundance were from comammox-*Nitrospira* (Palomo *et al*., 2016, Tatari *et al*., 2017). The strict AOB OTU found in this study was affiliated with oligotrophic *Nitrosomonas* cluster 6a which exhibit maximum growth rates at ammonia concentrations similar to the ones used for high and mid nitrogen amendments (Bollmann *et al*., 2011, Sedlacek *et al*., 2019). Despite this, the proportions of SSU reads mapping to *Nitrosomonas*-like populations in all nitrogen amendments were consistently low. Taxonomic classification of nitrogen cycling genes revealed metabolic potential for nitrification processes were confined to *Nitrospi*ra- and *Nitrosomonas*-like populations corroborating with assembled 16S rRNA gene sequences. Additionally, phylogeny of *amoA* sequences found in the metagenome indicated ammonia oxidation could be mediated by both *Nitrospira* and *Nitrosomonas*.

We assembled a total of nine nitrifier MAGs which included comammox-*Nitrospira* (n=4), *Nitrospira*-NOB-like (n=2), and *Nitrosomonas*-like (n=3) populations. Three of the four comammox MAGs assembled were identified as clade A2 based on phylogenetic analyses of hydroxylamine dehydrogenase (*hao*) which has previously been shown to dominate drinking water biofilters along with comammox clade B (Palomo *et al*., 2019). The remaining comammox MAG assembled from biofilter media in this study was affiliated with clade A1 based on *hao* gene phylogeny, which while atypical for drinking water biofilters is consistent with previously published metagenome from the Ann Arbor drinking water filters (Pinto *et al*., 2016). Similar coexistence of clade A1 and A2 comammox bacteria with canonical nitrifiers has been observed in tertiary rotating biological contactors treating municipal wastewater with low ammonium concentrations (Spasov *et al*., 2020). However, phylogenomic placement of clade A sub-groups in this study separated the comammox MAGs into distinct clusters associated with freshwater (Bin_13, clade A1), groundwater biofilters (Bin_49_4, clade A2) and tap water (Bin_260 and Bin_49_2_2, clade A2). Maintenance of high functional redundancy for the complete ammonia oxidation pathway may rely on coexisting comammox populations avoiding direct competition through distinct physiological niches. Additionally, the innocula were sourced from low substrate conditions which may also allow for the coexistence of multiple comammox populations. Strict AOB MAGs obtained in this study associated with low ammonia adapted *Nitrosomonas* cluster 6a (Koops *et al*., 2006) which is consistent with the inocula source being an oligotrophic environment (i.e., DWTPs). Furthermore, close relatives of *Nitrospira*-NOB MAGs obtained in this study originated from a tap water source where *Nitrospira*-NOB also coexisted with strict AOB and comammox bacteria under oligotrophic conditions (Wang *et al*., 2017). Our findings, consistent with previous studies, confirm the nitrifier community encompassed multiple populations capable of single and two-step nitrification within a single system with Nitrospira as the dominant nitrifier. However, the mechanism behind high abundances of Nitrospira-NOB in biofilters is not yet completely understood. Further, assessment of metabolic versality revealed initiation of nitrification through urea degradation was possible by all three nitrifying guilds. Though ureolytic activity is a widespread trait among cultured comammox-*Nitrospira* representatives and curated MAGs, the capability is confined to only some *Nitrospira*-NOB and *Nitrosomonas* species (Koch *et al*., 2015, Sedlacek *et al*., 2019). Here in particular, this a would allow *Nitrospira*-NOB to play a role in nitrite production in urea microcosms by crossing feeding ammonia from urea degradation to strict AOB, a mutualistic strategy which may not be active in ammonia amended microcosms.

### Comammox bacterial abundance increased irrespective of nitrogen source or loading but may compete with NOB depending on nitrogen source type

We tested the impact of nitrogen source and loading rates on temporal dynamics of a mixed nitrifying community to determine whether comammox bacteria are outcompeted at higher concentrations and/or favored in urea amendments due to their ureolytic activity. qPCR-based abundance tracking revealed comammox bacteria demonstrated a preferential enrichment over strict AOB in the nitrogen amendments irrespective of nitrogen source or availability. Additionally, strict AOB abundance did not exhibit any significant difference across the nitrogen amendment types. This is in contrast to previous work in soil microcosms where AOB abundance increased in response to high ammonia amendments (He *et al*., 2021). However, strict AOB populations in these soil microcosms were primarily *Nitrosospira* compared to oligotrophic *Nitrosomonas* cluster 6a which were the primary AOB in this study. Here, both comammox bacteria and strict AOB demonstrated increased abundance in all amendments during the earlier weeks of the experiment. Ultimately, while comammox bacteria were enriched over time our findings demonstrated this increased abundance was not associated with a decrease in the abundance of strict AOB in any of the nitrogen amendments. This suggests a lack of direct competition between the two comammox and strict AOB which could be attributable to the two ammonia oxidizers occupying separate nitrogen availability niches (Martens-Habbena *et al*., 2009, Kits *et al*., 2017). Stable abundances of strict AOB compared to enrichment of comammox could be due to a combination of factors ranging from (1) higher abundances of comammox bacteria in the inocula and (2) significantly higher biomass yields per mole of ammonia oxidizers for comammox bacteria compared to AOB (Kits *et al*., 2017).

Clade A2 associated comammox bacterial MAGs were dominant in the inocula and over the course of the experiment showed increased abundance in all amendments. In contrast, comammox bacteria belonging to clade A1 were lower in abundance and did not demonstrate significant change over time in any amended microcosm. Though physiological differences between comammox bacteria clades/sub-clades have yet to be established, earlier studies of DWTP biofilters have observed higher abundances of clade B (Fowler *et al*., 2018) or alternatively both clades found at the same DWTP but within separate rapid sand filters, where clade B was more abundant in the secondary filters receiving lower ammonia concentrations (Poghosyan *et al*., 2020). In this study, the lack of clade A1 enrichment over the course of the experiment may also indicate distinct physiological niches within clades (i.e., subclade-level niche differentiation). Future research is necessary to develop a clearer understanding of physiological differences between comammox bacteria at the clade/sub-clade level. Since cultivability of comammox bacteria remains an ongoing challenge, integrating multiple ‘omics techniques (i.e., metatranscriptomics and metaproteomics) may be an appropriate strategy for examining ammonia utilization and the expressed metabolisms of multiple coexisting comammox bacteria populations alongside canonical nitrifiers.

The negative association between comammox bacteria and canonical NOB observed in ammonia amendments could be a result of nitrite limitation resulting from complete nitrification driven by comammox bacteria. The possibility of comammox bacteria being a source of leaked nitrite to Nitrospira-NOB seems unlikely in this case as this would likely form a positive association between the two. Nitrite limitation driven competition between comammox bacteria and NOB is supported by the fact the negative associations between the groups were stronger at medium (3.5 mg-N/l) and low (1.5 mg-N/l) nitrogen availability as compared to the high ammonia amendments (i.e., 14 mg-N/l). In contrast, there was no significant association between the abundance of comammox bacteria and Nitrospira-NOB in the urea amended systems irrespective of nitrogen loading. We hypothesize that this variable observations between ammonia and urea amended systems likely emerge from the extent of metabolic coupling between AOB and NOB and the resultant ability of comammox to outcompete NOB. Specifically, while the rate of nitrite availability for NOB in ammonia amended systems is largely dictated by ammonia oxidation activity of AOB it is likely that nitrite availability in urea amended systems would be dictated by a combination of both AOB activity and indirectly by NOB. In this case, the production of nitrite could be mediated by *Nitrospira*-NOB capable of ureolytic activity by crossing feeding ammonia to strict AOB who in turn provide nitrite at a rate at which *Nitrospira*-NOB. This tight coupling between AOB and NOB is supported by stronger and more significant correlation between AOB and NOB abundance in urea amended systems as compared to ammonia amended systems. Thus, it appears that while comammox bacteria may outcompete *Nitrospira*-NOB in systems where AOB abundances are low and nitrite availability is largely dictated by AOB activity, this competitive exclusion may be limited in scenarios with established AOB-NOB cross feeding via urea where nitrite availability is governed not only by AOB’s ammonia oxidation rate but also by NOB’s ureolytic activity. Since urea is used directly by urease-positive nitrifiers, variabilities in their affinities for the substrate would play a role in the outcome of competition in urea microcosms but was not assessed in this study.

Altogether, our study demonstrates that comammox bacteria will dominate over canonical nitrifiers in communities sourced from nitrogen limited environments irrespective of nitrogen source type or loading rate without directly competing with canonical AOB. Further, our study also indicates comammox bacteria and AOB may occupy independent niches in communities sources from low nitrogen environments. Interestingly, we see evidence of potential competitive exclusion of NOB by comammox bacteria governed by nitrogen source dependent metabolic coupling between AOB and NOB.

## Supporting information

List of Figures

## Data availability

Raw sequence reads, metagenome assembly, and MAGs are available on NCBI at Bioproject number PRJNA764197.

## Funding sources

This work was supported by NSF Graduate Research Fellowship and Cochrane Fellowship to KV and by NSF Award number: 1703089.

## Notes

### Competing Interest Statement

The authors have declared no competing interest.

