## Supplementary Material for "Comammox *Nitrospira* bacteria outnumber canonical nitrifiers irrespective of nitrogen source and availability"

**Table S1: Concentration of DNA extracted from GAC microcosms. Replicate biomass samples were taken weekly from each condition (n=6) for a total of 96 samples.**

| Time Point | Sample | DNA concentration (ng/µL) |
| --- | --- | --- |
| Week 0 | Inocula | 224 |
| Week 1 | 14A | 104 |
|  | 14A | 98.2 |
|  | 3.5A | 106 |
|  | 3.5A | 91.4 |
|  | 1.5A | 166 |
|  | 1.5A | 140 |
|  | 14U | 95.7 |
|  | 14U | 133 |
|  | 3.5U | 181 |
|  | 3.5U | 130 |
|  | 1.5U | 105 |
|  | 1.5U | 107 |
| Week 2 | 14A | 84.6 |
|  | 14A | 83.1 |
|  | 3.5A | 170 |
|  | 3.5A | 102 |
|  | 1.5A | 102 |
|  | 1.5A | 96.7 |
|  | 14U | 150 |
|  | 14U | 119 |
|  | 3.5U | 107 |
|  | 3.5U | 103 |
|  | 1.5U | 143 |
|  | 1.5U | 95.7 |
| Week 3 | 14A | 64.2 |
|  | 14A | 70.8 |
|  | 3.5A | 76.7 |
|  | 3.5A | 91 |
|  | 1.5A | 88.8 |
|  | 1.5A | 95.6 |
|  | 14U | 79.2 |
|  | 14U | 77.4 |
|  | 3.5U | 74.5 |
|  | 3.5U | 66.2 |
|  | 1.5U | 61.3 |
|  | 1.5U | 128 |
| Week 4 | 14A | 48.7 |
|  | 14A | 75 |
|  | 3.5A | 82.6 |
|  | 3.5A | 83.3 |
|  | 1.5A | 93 |
|  | 1.5A | 76.5 |
|  | 14U | 114 |
|  | 14U | 88.9 |
|  | 3.5U | 153 |
|  | 3.5U | 97 |
|  | 1.5U | 162 |
|  | 1.5U | 108 |
| Week 5 | 14A | 79.6 |
|  | 14A | 84.5 |
|  | 3.5A | 91.5 |
|  | 3.5A | 101 |
|  | 1.5A | 95.4 |
|  | 1.5A | 88.3 |
|  | 14U | 104 |
|  | 14U | 100 |
|  | 3.5U | 88.8 |
|  | 3.5U | 99.3 |
|  | 1.5U | 92.2 |
|  | 1.5U | 68.9 |
| Week 6 | 14A | 134 |
|  | 14A | 104 |
|  | 3.5A | 141 |
|  | 3.5A | 80.2 |
|  | 1.5A | 93 |
|  | 1.5A | 105 |
|  | 14U | 98.2 |
|  | 14U | 83.2 |
|  | 3.5U | 76.2 |
|  | 3.5U | 86.3 |
|  | 1.5U | 95.5 |
|  | 1.5U | 83.2 |
| Week 7 | 14A | 73.5 |
|  | 14A | 74.7 |
|  | 3.5A | 81.6 |
|  | 3.5A | 88.7 |
|  | 1.5A | 79.1 |
|  | 1.5A | 84.2 |
|  | 14U | 65.2 |
|  | 14U | 67.8 |
|  | 3.5U | 113 |
|  | 3.5U | 71.2 |
|  | 1.5U | 89.6 |
|  | 1.5U | 71.4 |
| Week 8 | 14A | 79.1 |
|  | 14A | 120 |
|  | 3.5A | 88.1 |
|  | 3.5A | 76.6 |
|  | 1.5A | 84 |
|  | 1.5A | 80.7 |
|  | 14U | 89.4 |
|  | 14U | 75.8 |
|  | 3.5U | 76.5 |
|  | 3.5U | 96.5 |
|  | 1.5U | 117 |
|  | 1.5U | 69.2 |

**Table S2: Primers used for quantification of comammox bacteria *amoB*, *Nitrospira* 16S rRNA, AOB 16S rRNA and total bacteria 16S rRNA.**

| **Target** | **Primer Set** | **Forward Primer sequence**  **(5’-3’)** | **Reverse Primer sequence**  **(5’-3’)** | **Final concentration in PCR reaction** | **Annealing Temp (°C) /Time (sec)** | **Amplicon length (bp)** | **Reference** |
| --- | --- | --- | --- | --- | --- | --- | --- |
| Comammox clade A amoB gene | cmx_  amoB 148F/485R | TGGTAYGAYACNGAATGGG | CCCGTGATRTCCATCCA | 0.5 μM F/R | 52/45 | 337 | (Cotto et al., 2020) |
| Comammox clade A amoB gene | Mod_CMX_amoB 148F/485R | TGGTAYGAYACNSARTGGG | CCNGTGATRTCCATCCA | 0.5 μM F/R | 52/45 | 337 | This paper |
| Nitrospira 16S rRNA gene | Nspra675F - 746R | GCGGTGAAATGCGTAGAKATCG | TCAGCGTCAGRWAYGTTCCAGAG | 0.5 μM F/R | 58/30 | 93 | (Graham et al., 2007) |
| AOB 16S rRNA gene | CTO189FA/B/C*-RT1R | GGAGRAAAGCAGGGGATCG  GGAGGAAAGTAGGGGATCG | CGTCCTCTCAGACCARCTACTG | 0.4 μM F/R | 57/30 | 116 | (Hermansson and Lindgren 2001) |
| Total Bacteria 16S rRNA gene | F515 - R806 | GTGCCAGCMGCCGCGGTAA | GGACTACHVGGGTWTCTAAT | 0.2 μM F/0.4 μM R | 50/15 | 291 | (Caporaso et al., 2011) |

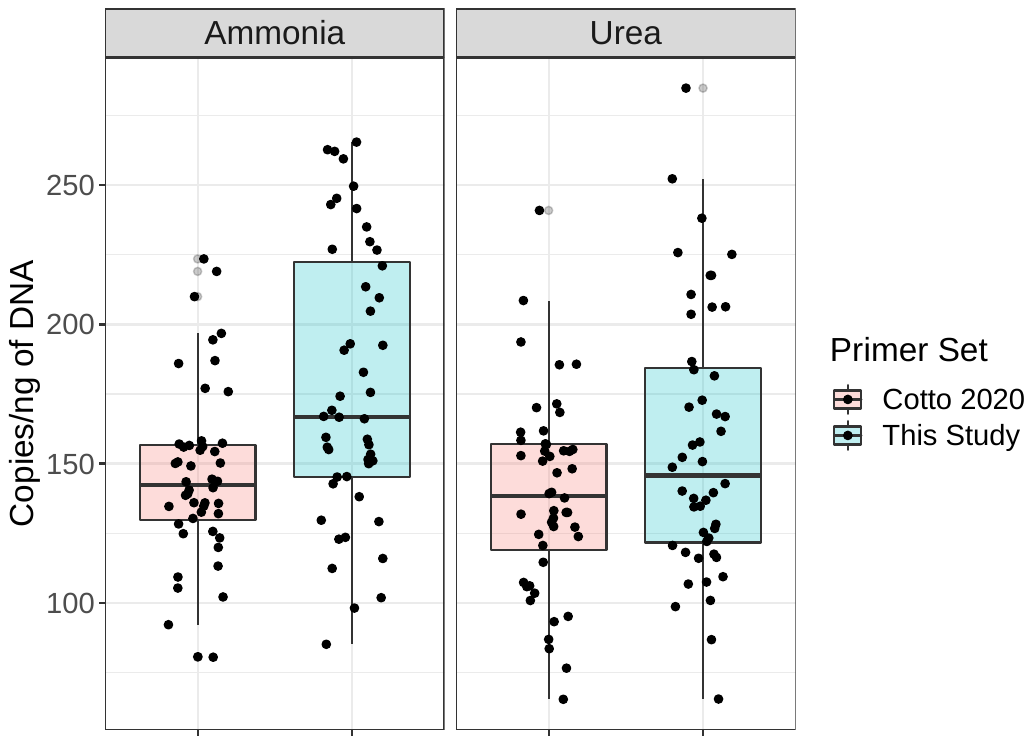

Figure S1: Comparison of qPCR-based quantification of comammox bacteria amoB gene using the primers developed by Cotto et al., 2020 (red) and the modified primers (blue) used in this study to inclusively capture amoB sequences of comammox clade A1 and A2.

**Table S3: Number of paired-end reads generated from two lanes of Illumina NovaSeq and the resulting number of paired-end reads after quality filtering using Fastp. 14A, 3.5A, and 1.A correspond to microcosms treated with 14 mg-N/l, 3/5 mg-N/l and 1/5 mg-N/l as ammonia while 14AU 3.5U, and 1.U correspond to microcosms treated with 14 mg-N/l, 3/5 mg-N/l and 1/5 mg-N/l as urea.**

| **Time Point** | **Treatment** | **Paired-end reads (millions)** | **Quality filtered paired-end reads (millions)** |
| --- | --- | --- | --- |
| Week 0 | Inocula | 68.27 | 67.83 |
| Week 4 | 14A | 34.87 | 34.67 |
|  | 3.5A | 31.62 | 31.43 |
|  | 1.5A | 30.46 | 30.28 |
|  | 14U | 34.67 | 34.46 |
|  | 3.5U | 29.65 | 29.44 |
|  | 1.5U | 40.77 | 40.57 |
| Week 8 | 14A | 46.79 | 46.53 |
|  | 3.5A | 32.39 | 32.19 |
|  | 1.5A | 39.36 | 39.05 |
|  | 14U | 38.47 | 38.25 |
|  | 3.5U | 32.95 | 32.75 |
|  | 1.5U | 34.86 | 34.63 |

**Table S4: Quast assembly statistics for Spades metagenomic assembly.**

| **Statistic (without reference)** | **Scaffold** |
| --- | --- |
| # contigs | 958730 |
| # contigs (>= 0 bp) | 2411090 |
| # contigs (>= 1000 bp) | 380914 |
| # contigs (>= 5000 bp) | 52805 |
| # contigs (>= 10000 bp) | 24369 |
| # contigs (>= 25000 bp) | 8531 |
| # contigs (>= 50000 bp) | 3430 |
| Largest contig | 2103352 |
| Total length | 2045559521 |
| Total length (>= 0 bp) | 2552730458 |
| Total length (>= 1000 bp) | 1649508864 |
| Total length (>= 5000 bp) | 1034572305 |
| Total length (>= 10000 bp) | 838880649 |
| Total length (>= 25000 bp) | 598410629 |
| Total length (>= 50000 bp) | 422206351 |
| N50 | 5199 |
| N75 | 1268 |
| L50 | 50492 |
| L75 | 277995 |
| GC (%) | 60.5 |
| **Mismatches** |  |
| # N's | 577321 |
| # N's per 100 kbp | 28.22 |

**Table S5: Accession number for reference genomes used for phylogenomic placement of nitrifying populations.**

| Genome | Functional Group | Accession # |
| --- | --- | --- |
| Candidatus Nitrospira defluvii | Nitrite oxidizing bacteria | GCA_000196815.1 |
| Nitrospira moscoviensis strain NSP M-1 | Nitrite oxidizing bacteria | GCA_001273775.1 |
| Nitrospira sp. bin75 | Nitrite oxidizing bacteria | GCA_002238765.1 |
| Nitrospira japonica NJ11 | Nitrite oxidizing bacteria | GCA_900169565.1 |
| Nitrospira cf moscoviensis SBR1015 isolate5 | Nitrite oxidizing bacteria | NZ_FJVN01000357.1 |
| Nitrospira sp UWLDO02 | Nitrite oxidizing bacteria | GCA_002254325.1 |
| Nitrospira sp CG24D | Nitrite oxidizing bacteria | GCA_002869855.2 |
| Nitrospira sp ST-bin5 | Nitrite oxidizing bacteria | GCA_002083555.1 |
| Nitrospira sp ND1 isolate | Nitrite oxidizing bacteria | GCA_900170025.1 |
| Nitrospira sp OLB3 | Nitrite oxidizing bacteria | GCA_001567445.1 |
| Ca Nitrospira inopinata | Comammox bacteria | GCA_001458695.1 |
| Ca Nitrospira nitrificans COMA2 | Comammox bacteria | GCA_001458775.1 |
| Ca Nitrospira nitrosa COMA1 | Comammox bacteria | GCA_001458735.1 |
| Nitrospira sp CG24A | Comammox bacteria | GCA_002869925.2 |
| Nitrospira sp CG24B | Comammox bacteria | GCA_002869845.2 |
| Nitrospira sp CG24C | Comammox bacteria | GCA_002869885.2 |
| Nitrospira sp CG24E | Comammox bacteria | GCA_002869895.2 |
| Nitrospira sp UWLDO01 | Comammox bacteria | GCA_002254365.1 |
| Nitrospira sp. Ga0074138 | Comammox bacteria | GCA_001464735.1 |
| Nitrospira sp isolate RCA | Comammox bacteria | GCA_005239465.1 |
| Nitrospira sp isolate RCB | Comammox bacteria | GCA_005239475.1 |
| Nitrospira sp SG-bin1 | Comammox bacteria | GCA_002083365.1 |
| Nitrospira sp SG-bin2 | Comammox bacteria | GCA_002083405.1 |
| Nitrospira sp ST-bin4 | Comammox bacteria | GCA_002083565.1 |
| Nitrospira sp UBA2082 | Comammox bacteria | GCA_002331335.1 |
| Nitrospira sp UBA2083 | Comammox bacteria | GCA_002331625.1 |
| Nitrospira sp UBA5698 | Comammox bacteria | GCA_002420115.1 |
| Nitrospira sp UBA5699 | Nitrite oxidizing bacteria | GCA_002420105.1 |
| Nitrospira sp UBA5702 | Comammox bacteria | GCA_002420045.1 |
| Nitrospira sp UBA6909 | Comammox bacteria | GCA_002451055.1 |
| M DeepCast 65m mx 150 | Comammox bacteria | requested |
| M DeepCast 50m m2 151 | Comammox bacteria | requested |
| Nitrosomonas communis strain Nm2 | Ammonia oxidizing bacteria | GCA_001007935.1 |
| Nitrosomonas cryotolerans ATCC 49181 | Ammonia oxidizing bacteria | GCA_900143275.1 |
| Nitrosomonas europaea ATCC 19718 | Ammonia oxidizing bacteria | GCA_000009145.1 |
| Nitrosomonas eutropha strain Nm14 | Ammonia oxidizing bacteria | GCA_900100815.1 |
| Nitrosomonas halophila strain Nm1 | Ammonia oxidizing bacteria | GCA_900107165.1 |
| Nitrosomonas nitrosa strain Nm146 | Ammonia oxidizing bacteria | GCA_900114795.1 |
| Nitrosomonas oligotropha strain Nm49 | Ammonia oxidizing bacteria | GCA_003050805.1 |
| Nitrosomonas sp. AL212 | Ammonia oxidizing bacteria | GCA_000175095.2 |
| Nitrosomonas sp. Is79A3 | Ammonia oxidizing bacteria | GCA_000219585.1 |
| Nitrosomonas ureae strain Nm10 | Ammonia oxidizing bacteria | GCA_001455205.1 |
| Nitrosospira briensis C 128 | Ammonia oxidizing bacteria | GCA_000619905.2 |
| Nitrosospira lacus strain APG3 | Ammonia oxidizing bacteria | GCA_000355765.4 |
| Nitrosospira multiformis ATCC 25196 | Ammonia oxidizing bacteria | GCA_000196355.1 |

**Table S6: Statistics for 145 metagenome assembled genomes obtained in this study. See excel sheet Supplemental MAG Information.**

**Table S7: Statistics for nitrifying metagenome assembled genomes in this study*.* The completeness and redundancy of each MAG was determined using CheckM (version 1.07).**

| **Nitrifying MAGs** | **Bin_49_2_2** | **Bin_49_4** | **Bin_260** | **Bin_13** | **Bin_7_1** | **Bin_188** | **Bin_168** | **Bin_195** | **Bin_83** |
| --- | --- | --- | --- | --- | --- | --- | --- | --- | --- |
| Functional group | Comammox | Comammox | Comammox | Comammox | Nitrospira-NOB | Nitrospira-NOB | AOB | AOB | AOB |
| Completeness (%) | 96.76 | 91.31 | 88.63 | 89.09 | 38.04 | 48.25 | 83.79 | 50.6 | 99.04 |
| Redundancy (%) | 3.69 | 1.92 | 5 | 18.71 | 8.76 | 8.45 | 7.34 | 12.27 | 1.91 |
| Genome Size (Mbp) | 3.53 | 3.62 | 3.46 | 8.48 | 3.51 | 2.76 | 2.92 | 2.45 | 3.06 |
| N50 | 165773 | 64940 | 299090 | 10589 | 10951 | 37110 | 6076 | 6569 | 36991 |

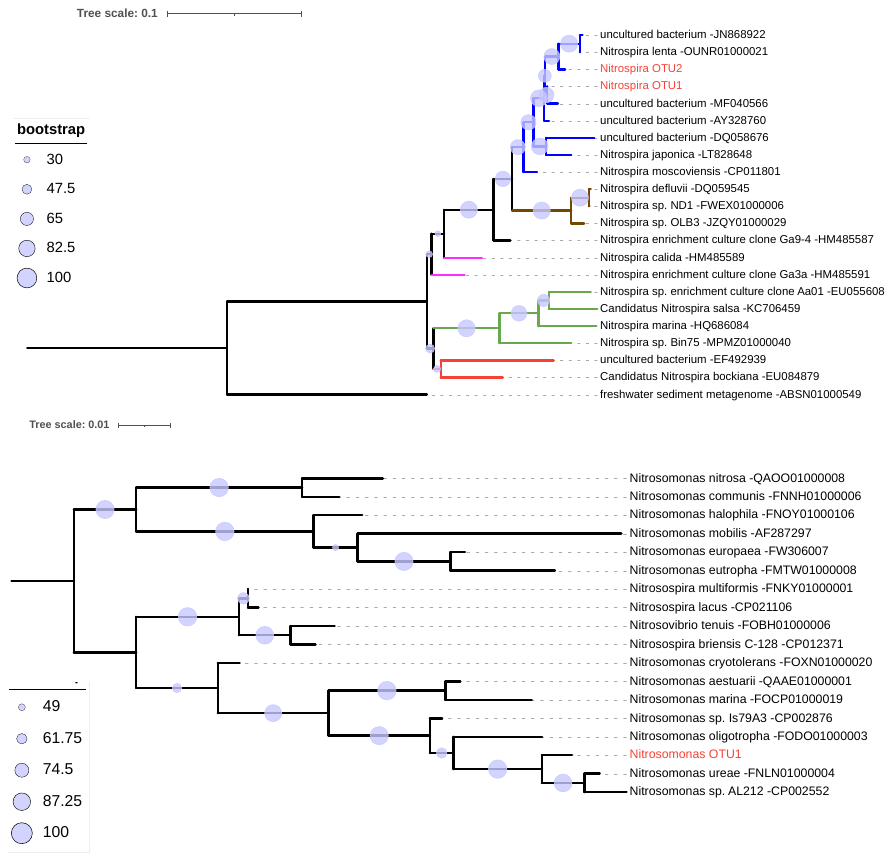

B

A

Figure S2: Maximum likelihood tree based on 16S rRNA gene sequences from Nitrospira references (black) and assembled OTUs in this study (red). Both Nitrospira OTUs clustered with sublineage II (blue branches). Brown, pink, green, and red branches represent sublineages I, VI, IV and V (A). Maximum likelihood tree based on 16S rRNA gene sequences from Nitrosomonadacae and the assembled OTU from this study (red) (B).

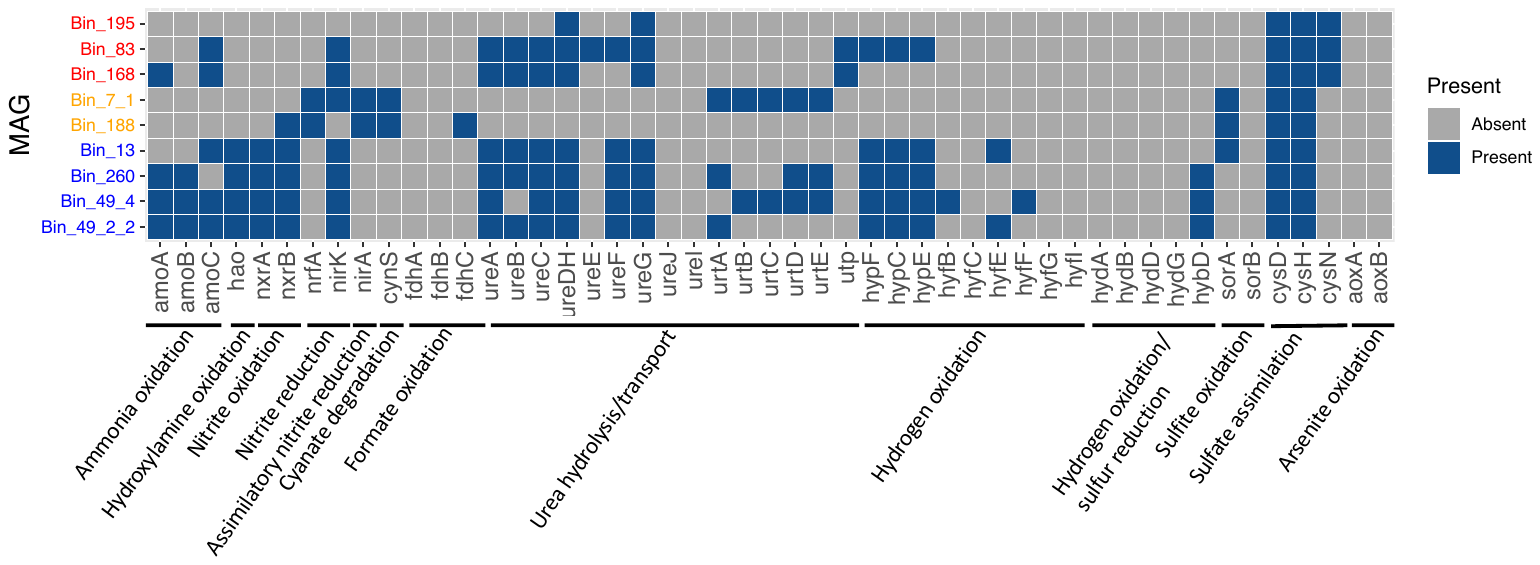

Figure S3: Presence/absence of genes involved in the nitrogen metabolism in MAGs identified as nitrifiers based on taxonomic classification. The gene annotation was performed against the KEGG database using kofamscan (version). Nitrosomonas, Nitrospira-NOB and comammox bacteria MAGs are colored red, orange and blue, respectively.

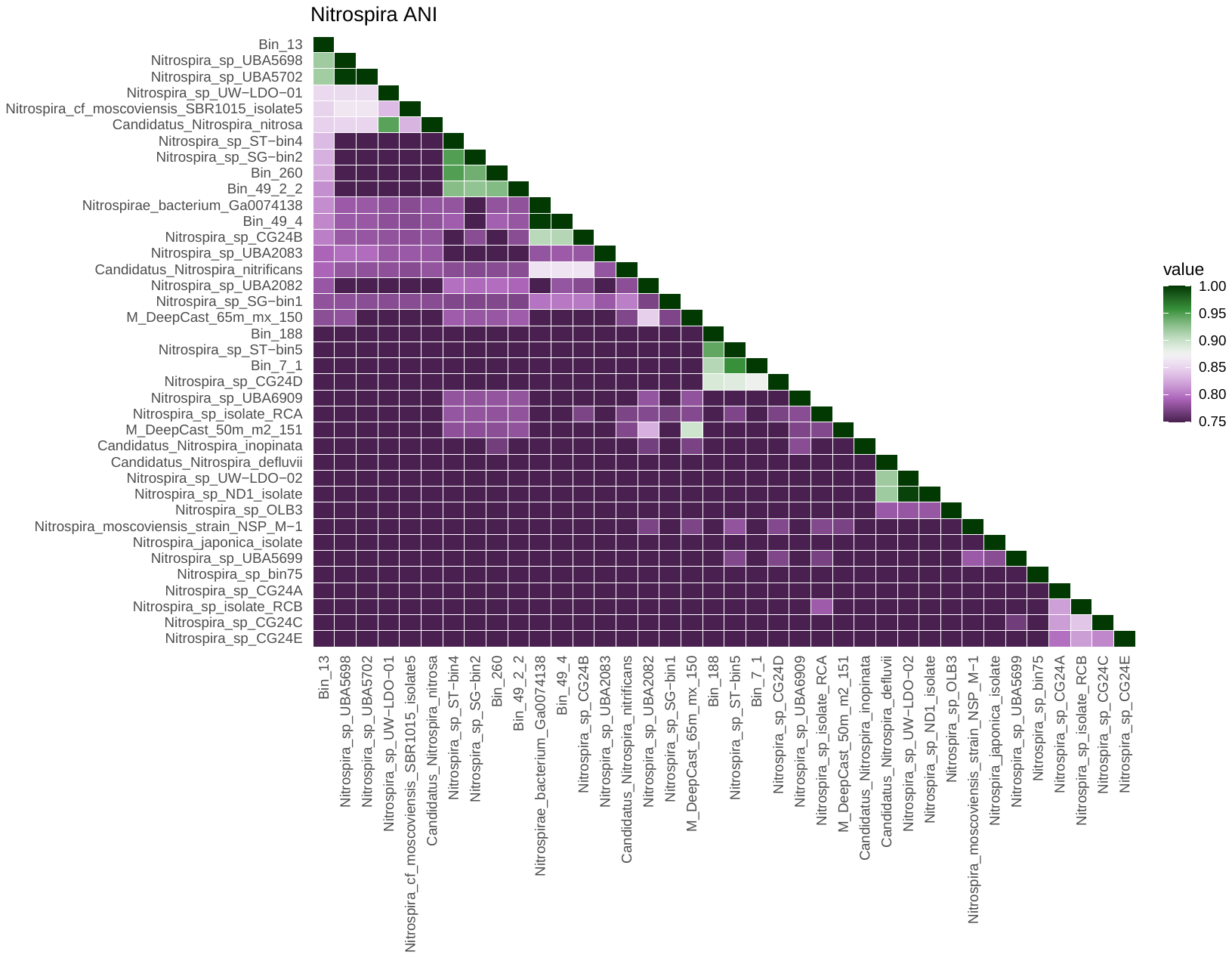

Figure S4: Heatmap indicating the average nucleotide identity (ANI) between comammox bacterial MAGs detected in this study with 38 canonical Nitrospira-NOB and comammox bacteria genomes/MAGs estimated using fastANI (version). Most comammox MAGs obtained in this study had the highest similarities to references obtained from biofilters and drinking water systems. fastANI does not report ANI values for genome pairs with less than 75% ANI, thus dark purple here represents an ANI of 75% or less.

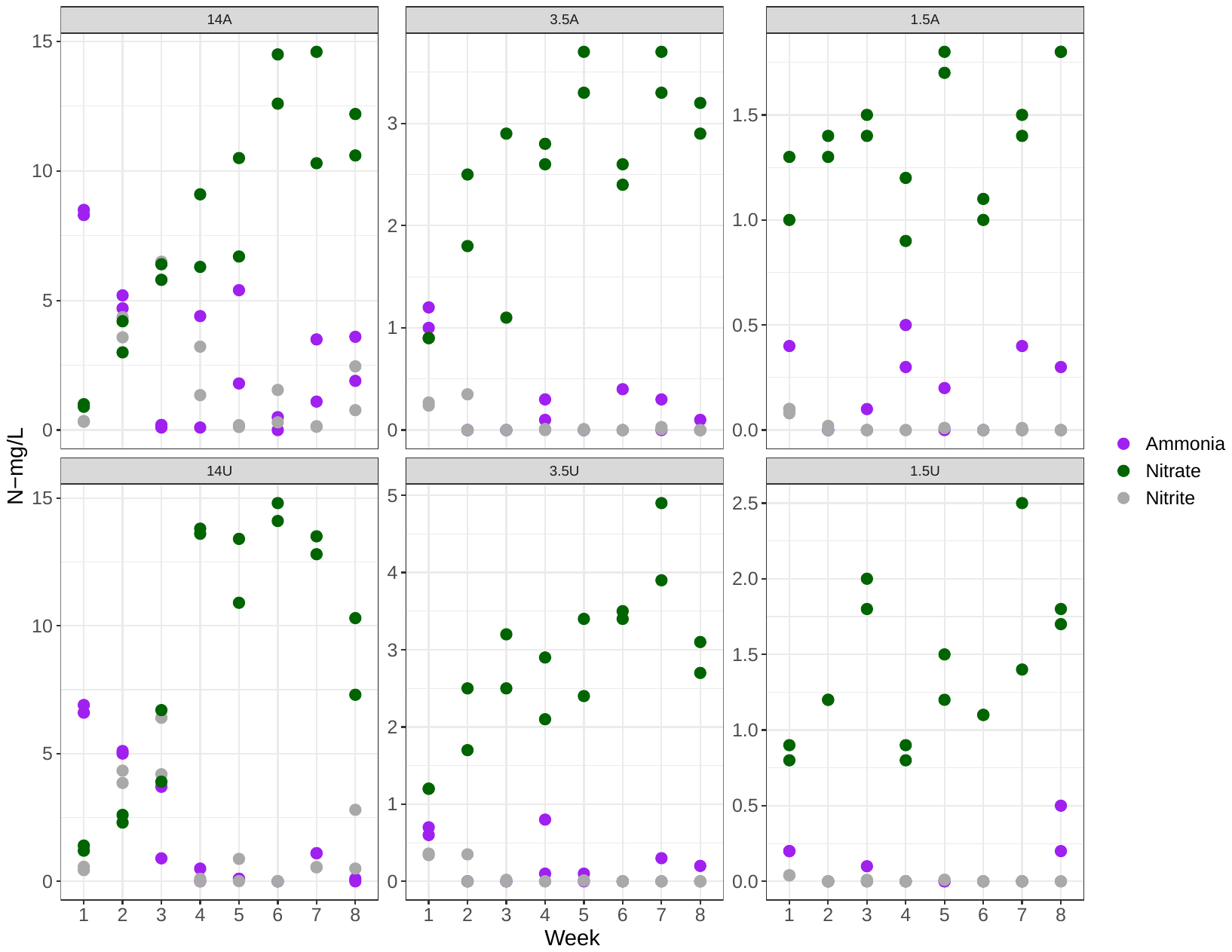

Figure S5: Weekly ammonia, nitrite, and nitrate concentrations measured in microcosms from each nitrogen amendment. Data

points are the average concentration of biological replicate microcosms with bars for standard deviation.

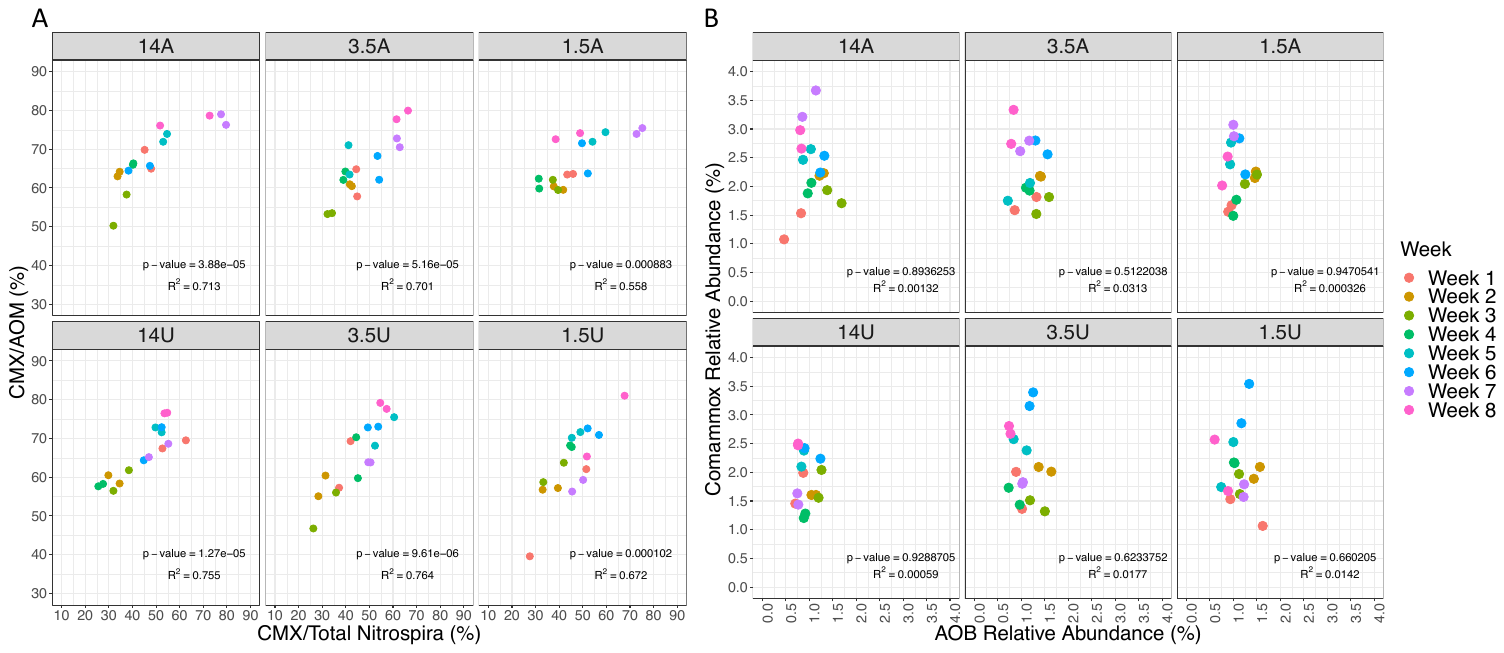

Figure S6: A) Relationship between the abundance of comammox bacteria as a proportion of ammonia oxidizing microorganisms (CMX:AOM) and comammox bacteria as a proportion of Nitrospira bacteria (CMX:Total Nitrospira). All nitrogen amendments exhibit a strong linear relationship. B) There was no significant association between changes in comammox bacteria concentration and that of AOB as a proportion of total bacteria.
