## Supplementary material for "Comammox *Nitrospira* bacteria outnumber canonical nitrifiers irrespective of nitrogen source and availability": List of Figures

**Figure 1**: A) Proportion of metagenomic reads mapping to the SILVA SSU NR99 database (version 138.1) from the inocula and treated samples at weeks four and eight. The community primarily consisted of Gammaproteobacteria, Alphaproteobacteria and Nitrospirota. Proteobacteria is broken down by classes, Alphaproteobacteria and Gammaproteobacteria, though a small portion of Proteobacteria reads could not be classified further. B) Taxonomic classification of genes for nitrogen biotransformation present in the metagenome at the phyla level with Proteobacteria presented by classes, Alphaproteobacteria and Gammaproteobacteria. C and D) Phylogenetic placement of amoA-pmoA like sequences (C), and nxrA sequences (D) detected in the metagenomes. Both maximum likelihood trees were constructed based alignments of protein sequences of the respective genes. Sequences identified in this study are colored according to their phylogenetic placement (red, green, blue and orange) while references are black. AOB = ammonia oxidizing bacteria, MOB = methane oxidizing bacteria.

**Figure 2**: Phylogenomic tree for Nitrospira MAGs (blue and green) obtained in this study and 32 reference genomes (black). Blue label = comammox, green label = NOB. Branch colors represent different Nitrospira sublineages. Two Leptospirillum reference genomes were used as the outgroup for maximum likelihood tree construction. B) Maximum likelihood tree based on ammonia monooxygenase subunit A (amoA) sequences from comammox-Nitrospira. C) Maximum likelihood tree based on hydroxylamine oxidoreductase (hao) sequences of comammox-Nitrospira. For B and C blue labels represent amoA/hao gene sequences found in comammox MAGs from this study, while black labels are reference sequences. Environment of origin is denoted with colored squares to the left of each tree. amoA and hao protein sequences from Nitrosomonas europaea and Nitrosomonas oligotropha were used as the outgroup for comammox trees in B and C, respectively. D) Phylogenomic tree for strict AOB MAGs (yellow) obtained in this study and 10 Nitrosomonas reference genomes (black). Three Nitrosospira genomes were used as the outgroup.

**Figure 3**: **Figure 3**: A) The relative abundance of comammox bacteria (blue) and strict AOB (red) as a proportion of all ammonia oxidizing microorganisms calculated using copy number of 16S rRNA and amoB genes of strict AOB and comammox bacteria, respectively and dividing by the combined copy number to represent total AOM for each time point. Data points averaged from triplicate qPCR analyses of samples across biological duplicate microcosms. B) Relative abundance of comammox bacteria (blue), Nitrospira-NOB (orange) and strict AOB (red) as a proportion of total bacteria using the ratio of copy number of the respective nitrifier genes to copy number of total bacteria averaged between duplicate samples. Panels display relative abundances of the nitrifiers subject to varying nitrogen amendments where data points represent the average of biological replicates (qPCR performed in triplicate) and error bars for standard deviation.

**Figure 4:** Reads per kilobase million (RPKM) calculated for all MAGs identifying as comammox bacteria (blue), Nitrospira-NOB (orange) and strict AOB (red) at selected time points.

**Figure 5**: A) Significant positive correlation between changes in AOB concentration and that of Nitrospira-NOB as a proportion of total bacteria were found in most treatments expect low ammonia (1.5A). B) Negative associations between changes in comammox bacteria concentration and that of Nitrospira-NOB as a proportion of total bacteria existed in ammonia amendments with statistically significance detected in 3.5A and 1.5A while no association existed in urea amendments.
